# Eye movements support behavioral pattern completion

**DOI:** 10.1101/764084

**Authors:** Jordana S. Wynn, Jennifer D. Ryan, Bradley R. Buchsbaum

## Abstract

The ability to recall a detailed event from a simple reminder is supported by *pattern completion*, a cognitive operation performed by the hippocampus wherein existing mnemonic representations are retrieved from incomplete input. In behavioral studies, pattern completion is often inferred through the false endorsement of lure (i.e., similar) items as old. However, evidence that such a response is due to the specific retrieval of a similar, previously encoded item is severely lacking. We used eye movement (EM) monitoring during a partial-cue recognition memory task to index reinstatement of lure images behaviorally via the recapitulation of encoding-related EMs or, *gaze reinstatement*. Participants reinstated encoding-related EMs following degraded retrieval cues and this reinstatement was negatively correlated with accuracy for lure images, suggesting that retrieval of existing representations (i.e., pattern completion) underlies lure false alarms. Our findings provide novel evidence linking gaze reinstatement and pattern completion and advance a functional role for EMs in memory retrieval.

## Introduction

Memory is often a kind of guessing game: we receive fragmentary, noisy, or degraded information from the sensory environment and then must “fill in” the blanks. Seeing a blurry face in the distance or hearing a note from a familiar song can suddenly lead to recognition of a close friend or the incessant humming of a tune. This remarkable ability is thought to rely on a neurocomputational process called “pattern completion” (for review, see Rolls, 2013; Yassa & Stark, 2011). *Pattern completion* refers to the retrieval of a complete memory representation from partial or degraded input, while the complementary process of *pattern separation* refers to the transformation of similar inputs into distinct, non-overlapping memory traces (Marr, 1971; McClelland, O’Reilly, & McNaughton, 1995; for review, see Hunsaker & Kesner, 2013). Both processes have been attributed to distinct subregions in the hippocampus, and contribute to its broader role in the encoding and retrieval of the relations among stimulus or event features (Cohen & Eichenbaum, 1993; Eichenbaum & Cohen, 2001).

Although pattern separation and pattern completion describe neurocomputational processes, they are often inferred from behavioral memory responses. That is, when a lure (i.e., similar) item (e.g., two different pictures of apples) is correctly identified as new, pattern separation is thought to have occurred; whereas when a lure item is incorrectly identified as old, pattern completion is thought to have occurred (Bakker, Kirwan, Miller, & Stark, 2008; Clelland et al., 2009; Stark, Yassa, Lacy, & Stark, 2013; Toner, Pirogovsky, Kirwan, & Gilbert, 2009). While previous work has often relied on such responses to index pattern completion, this approach is indirect and rests on untested assumptions; namely, that false alarms to lure stimuli are the result of retrieval of the originally-encoded item (see also, Cowell, Barense, & Sadil, 2019; Molitor, Ko, Hussey, & Ally, 2014). Presumably, the operation of pattern completion should entail the retrieval of individual stimulus features absent from the test probe as well as the relations between those features and features present in the test probe (see also, Hannula, Ryan, Tranel, & Cohen, 2007). However, that retrieval of such a detailed relational representation has occurred cannot be assumed from a simple behavioral response. Thus, to evaluate this claim more directly, the present study used eye movement (EM) monitoring to measure the overlap between participant- and stimulus-specific gaze patterns during encoding and retrieval (visualization) of degraded lure stimuli as a predictor of memory performance. Whereas behavioral responses reflect the outcome of an underlying retrieval process, EM-based reinstatement has been linked to the online maintenance and retrieval of relational information (for review, see Wynn, Shen, & Ryan, 2019), making it a powerful tool to assess reactivation.

Eye movements provide a unique window into cognitive processes as they unfold in time (for review, see Hannula, Althoff, Warren, Riggs, & Cohen, 2010). Research using EM monitoring suggests that EMs are involved in the binding of stimulus features into cohesive memory representations during encoding, and the retrieval of those features and the relations among them during retrieval (for review, see Hannula et al., 2010, 2007; Wynn, Shen, et al., 2019). For example, whereas fixed viewing impairs memory (Armson, Ryan, & Levine, 2019; Bochynska & Laeng, 2015; Henderson, Williams, & Falk, 2005; Johansson, Holsanova, Dewhurst, & Holmqvist, 2012; Johansson & Johansson, 2013), spontaneous gaze shifts to regions corresponding with previously encoded (i.e., viewed) image features has been shown to facilitate reactivation of those features and the relations among them (Noton & Stark, 1971a, 1971b; for review, see Wynn, Shen, et al., 2019). Reinstatement of encoding-related EMs, or *gaze reinstatement,* during memory maintenance (Olsen, Chiew, Buchsbaum, & Ryan, 2014; Wynn, Olsen, Binns, Buchsbaum, & Ryan, 2018) and retrieval (Damiano & Walther, 2019; Holm & Mäntylä, 2007; Laeng & Teodorescu, 2002; Wynn et al., 2016) has been associated with mnemonic performance across a variety of tasks. Even in the absence of visual input, humans spontaneously direct their gaze to image regions previously inspected during encoding (i.e., “looking at nothing”), and this gaze reinstatement has been correlated with explicit measures of memory. (Bochynska & Laeng, 2015; Bone et al., 2018; Ferreira, Apel, & Henderson, 2008; Foulsham & Kingstone, 2013; Johansson & Johansson, 2013; Laeng, Bloem, D’Ascenzo, & Tommasi, 2014; Laeng & Teodorescu, 2002; Scholz, Mehlhorn, & Krems, 2016; Wynn et al., 2016; for review, see Wynn, Shen, et al., 2019).

Extending this work, recent neuroimaging findings suggest that gaze reinstatement may rely on the same neural mechanisms that support memory retrieval. For example, a recent study from Bone et al. (2018) found that participants reinstated encoding-related EMs during stimulus-free visualization, and this reinstatement was positively correlated with whole-brain neural reactivation (i.e., similarity between image-specific patterns of brain activity evoked during perception and imagery), which in turn was correlated with objective (change detection performance) and subjective (vividness ratings) measures of memory. Gaze reinstatement that differentiates hits from misses for configurally similar scene images has also been correlated with activity in the hippocampus (Ryals, Wang, Polnaszek, & Voss, 2015), supporting the existence of a functional link between EMs and hippocampally-mediated relational memory processes (also see, Bicanski & Burgess, 2019; Kragel et al., 2019; Liu, Shen, Olsen, & Ryan, 2017; Nau, Julian, & Doeller, 2018; Ryan et al., 2018; Shen, Bezgin, Selvam, McIntosh, & Ryan, 2016; Voss, Bridge, Cohen, & Walker, 2017; for review, see Hannula, Ryan, & Warren, 2017). Thus, given that EMs and memory retrieval, and the neural networks underlying them, are intimately related, the present study used EM monitoring to assess the online reactivation of previously encoded stimuli during mnemonic discrimination of lure images via gaze reinstatement.

From a computational standpoint, pattern completion involves the reactivation of a total memory trace from a partial or degraded stimulus that serves as an input pattern to an auto-associative network (for review, see Hunsaker & Kesner, 2013; Santoro, 2013). Thus, to capture pattern completion as it has been defined in computational models, we used memory cues that had been systematically degraded by randomly removing chunks of information from the recognition probes. After an image-encoding phase, participants were briefly (<750ms) presented with old and lure test images that were manipulated such that a proportion of the image (0-80%) was occluded. Before making an old/new recognition response, participants were instructed to visualize the presented image while looking at a blank screen. Importantly, the lack of visual input during this post-test interval allowed us to isolate the effect of memory on retrieval-related EMs from the effects of the visual properties of the test probe.

To determine whether the degraded test probe elicited retrieval of the corresponding (same or similar) encoded stimulus, we computed the overlap between encoding- and retrieval-related gaze patterns. If false alarms to lure stimuli are indeed related to retrieval of the similar item via pattern completion, this should be reflected in the accompanying EMs, which should be directed towards screen regions previously visited during encoding of the originally presented stimulus. Accordingly, we hypothesized that similarity between encoding- and retrieval-related gaze patterns during the post-test interval should be significantly greater than chance even when images are substantially degraded, indicating that the corresponding encoded representation has been reactivated. Moreover, based on previous evidence of gaze reinstatement and behavioral pattern completion, we predicted that reinstatement of encoding-related EMs during partially cued retrieval would be positively correlated with accuracy for old images and negatively correlated with accuracy for lure images.

To further examine the content and specificity of retrieved representations, we computed retrieval-related reactivation of the degraded test probe image (i.e., “probe reinstatement”) along with two separate measures of encoding-retrieval EM similarity to measure both the idiosyncratic reactivation of stimulus-specific features and relations (i.e., “gaze reinstatement”) and the participant-invariant reactivation of non-specific image features and relations (i.e., “image reinstatement”). We then compared the strength of each measure in predicting retrieval-related EMs and accuracy across the retrieval interval. Critically, by computing multiple measures of EM-based reinstatement, we were able to not only deduce the occurrence of pattern completion, that is, that a specific previously-encoded item has been retrieved, but also the nature of the pattern being completed, and specifically whether it comprises the stimulus itself (image reinstatement) or the stimulus *and* the operation by which it was encoded (gaze reinstatement).

## Methods

### Participants

Participants were 64 young adults (43 female) aged 19-35 (*M*= 23.66, *SD*= 3.85) with normal or corrected-to-normal vision who were recruited through the Rotman Research Institute’s participant database. All participants provided informed consent before participating in the experiment in accordance with the ethical guidelines of the Rotman Research Institute and were compensated at a rate of $10/hr for their participation. Seven participants were excluded from analysis on the basis of missing data (*n* =2), average performance (corrected recognition) lower than 2.5 SD from the mean (*n* =2), average gaze reinstatement greater than 3.5 SD from the mean^1^ (*n* =1), and failure to follow instructions^2^ (*n* =2). Data from the remaining 57 participants were analyzed.

### Apparatus

Stimuli were presented on a on a 1920 x 1080 resolution, 19 inch Dell M991 monitor. Monocular EMs were recorded using a head-mounted EyeLink II eyetracking system at 500 Hz sampling rate (SR Research Ltd., Mississauga, Canada). Eye movement calibration was accomplished using a 9-point calibration procedure, which was performed prior to the experiment. Drift correction (>5°) was performed between trials. Saccades and blinks were defined by EyeLink as saccades greater than 0.5° of visual angle and the period in which saccade signal was missing for three or more consecutive samples, respectively. All remaining samples were classified as fixations.

### Stimuli

Stimuli consisted of 240 images, or 120 sets (A-B) of unique, but similar 800 x 600 pixel images displayed against a black background. During the test phase, image duration (250, 500, 750ms) and degradation (0%, 20%, 40%, 60%, 80%) were manipulated. Images were degraded by randomly placing 100 x 100 pixel grey squares over the image, see Fig 1. Images were randomly assigned to one of 4 study/test blocks and counterbalanced across duration, degradation, and probe type (old, new). Of the 30 images presented during each study block, participants viewed 15 again as test probes (“old”). Lures from the alternate set of similar images were presented as test probes for the remaining 15 studied images.

**Fig 1.**
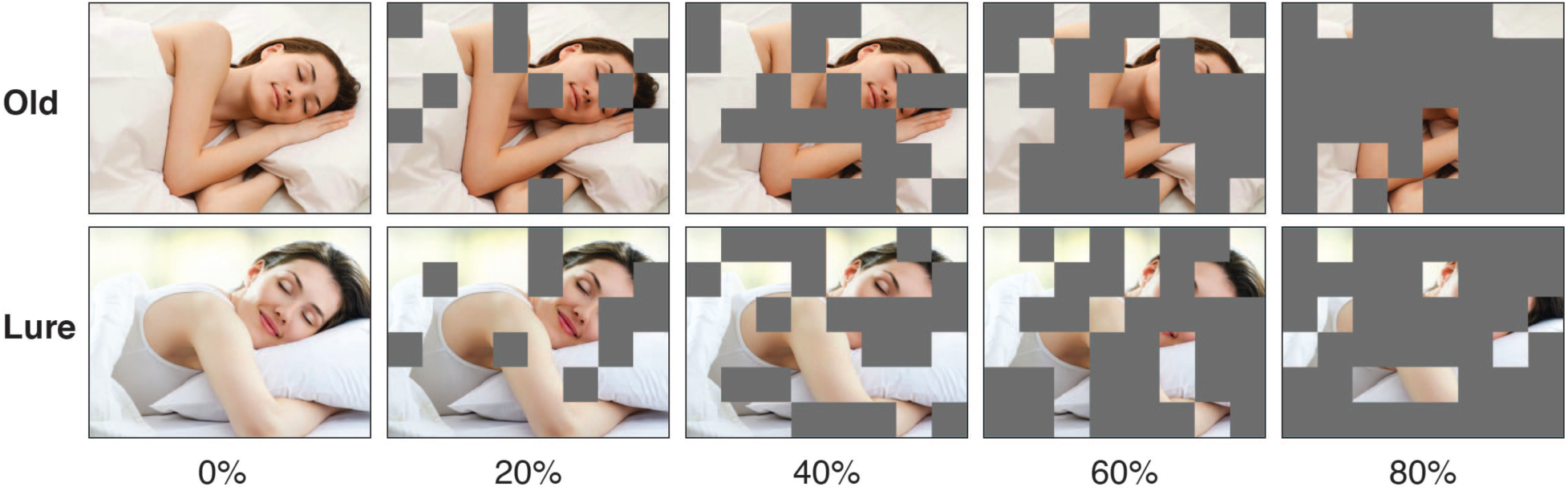
Examples of old and lure stimulus images at all levels of image degradation.

### Procedure

Before the start of the experiment, participants completed 2 short (6 trials) study-test practice blocks with novel images to familiarize themselves with the paradigm. Participants subsequently completed 4 blocks of a modified recognition memory test (Fig 2), with each block containing a novel set of images. During each study block, participants were instructed to view and memorize a set of 30 images, which were presented 4 times each. Image presentation order was randomized within each repetition. On each trial, participants were presented with a single image for 3s. A 2s fixation cross was presented between trials, during which the experimenter was able to perform online drift correction if necessary. Following the study block, participants were tested on their memory for the studied images during a test block. On each test trial, participants were presented with an old (presented at study), or new (lure: similar, but not identical to an image presented at study) image for 250, 500, or 750ms, followed by a 50ms mask. The purpose of the mask was to prevent sensory persistence from influencing viewing during the post-test interval. Image presentation was manipulated such that each image could be presented either in full (0% degradation), or with 20%, 40%, 60%, or 80% degradation (Fig 1). Participants were told to ignore the grey squares when making a response (i.e., to base their response on the visible portions of the underlying image). Following the mask, participants were presented with a grey square the same size as the study and test images (800 x 600 pixels) against a black background for 3s, during which they were instructed to visualize the presented test image. After this post-test interval, participants were given 3s to indicate whether the presented test image was “old” or “new” via key press. Participants were instructed to respond “old” only if the test image was exactly the same as an image presented during study, and to respond “new” to all other images (i.e., lures). Test trials were separated by a variable length (2 - 6s) fixation cross, during which online drift correction was performed at the discretion of the experimenter.

**Fig 2.**
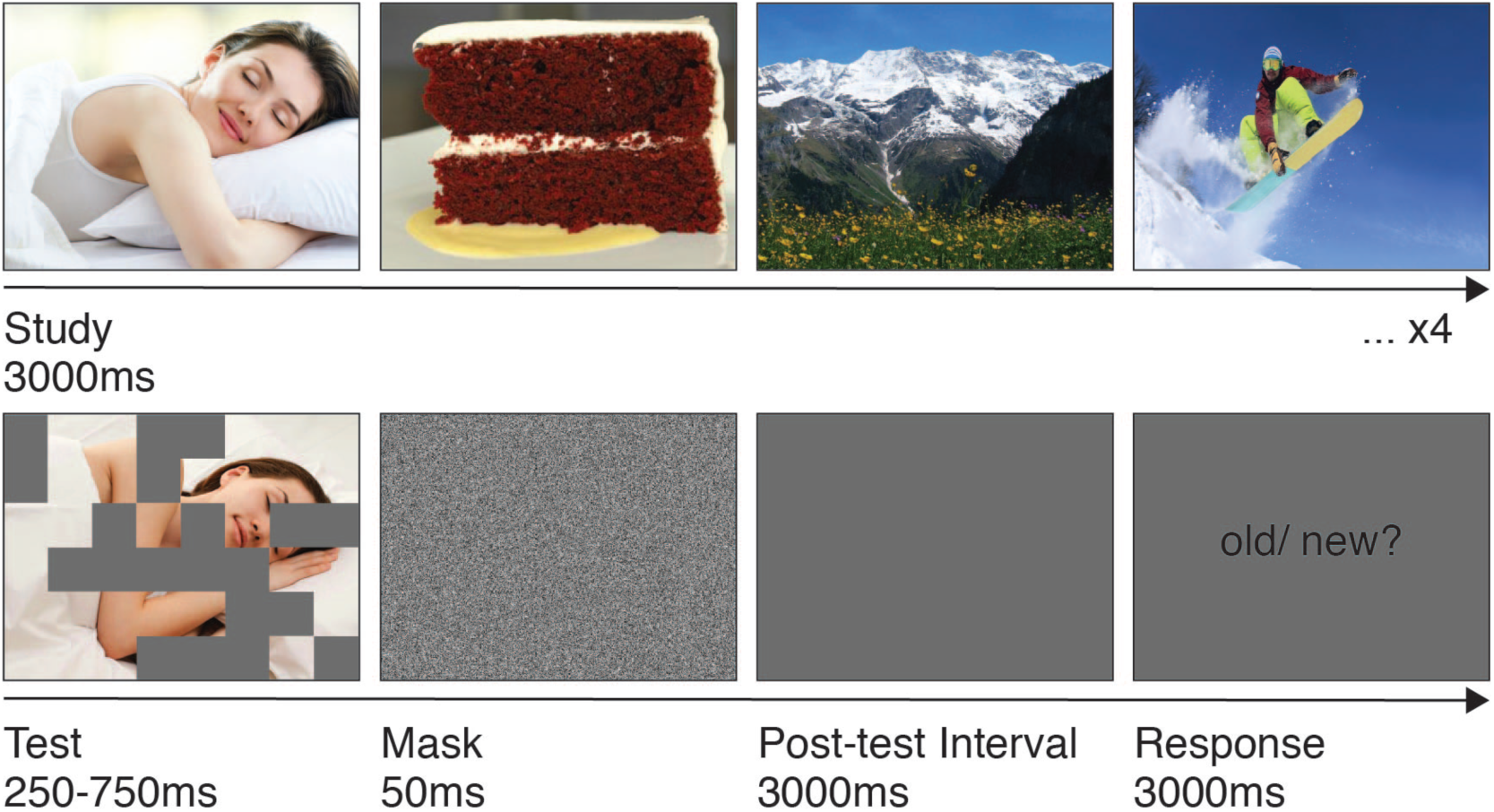
Experimental procedure. During the study period, each image was presented for 3000ms. A 2000ms fixation cross (not shown here) appeared between trials, to allow for online drift correction. Study images were presented 4 times each. During the test period, each test probe was presented for 250, 500, or 750ms, either in full (0% degradation), or with 20%, 40%, 60%, or 80% of the image obscured by 100 x 100pixel grey squares. Following the test probe, a visual mask was presented for 50ms, after which a grey square was presented for 3000ms, during which participants were instructed to visualize the presented image. Finally, participants were given 3000ms to indicate whether the presented image was “old” (presented at study) or “new” (i.e., lure: similar, but not identical to an image presented at study). A 2000-6000ms fixation cross (not shown here) appeared between trials, to allow for online drift correction. Note that all images and the grey post-test interval square were presented at 800 x 600pixels on a black background; the images have been expanded here for visualization.

### Eye movement analyses

In the present study, we computed three separate measures of EM-based reinstatement (Fig 3-4) using the R eyesim package (https://github.com/bbuchsbaum/eyesim). While we were primarily interested in EM-based reinstatement of information from long term memory (LTM), it is possible that EMs during the post-test interval reinstate salient regions of the just-presented test probe image. Therefore, in order to distinguish between reinstatement that is guided by memory for the encoded image from reinstatement guided by memory for the just-presented test probe, we computed a measure of *probe reinstatement,* reflecting the similarity between the EM patterns of *all* participants viewing the test probe (i.e., the visible portions of the test image weighted by the EM pattern of all participants encoding the same image during the study period), and a single participant subsequently retrieving that image (or the alternate lure image) during the 3s stimulus-free post-test interval.

**Fig 3.**
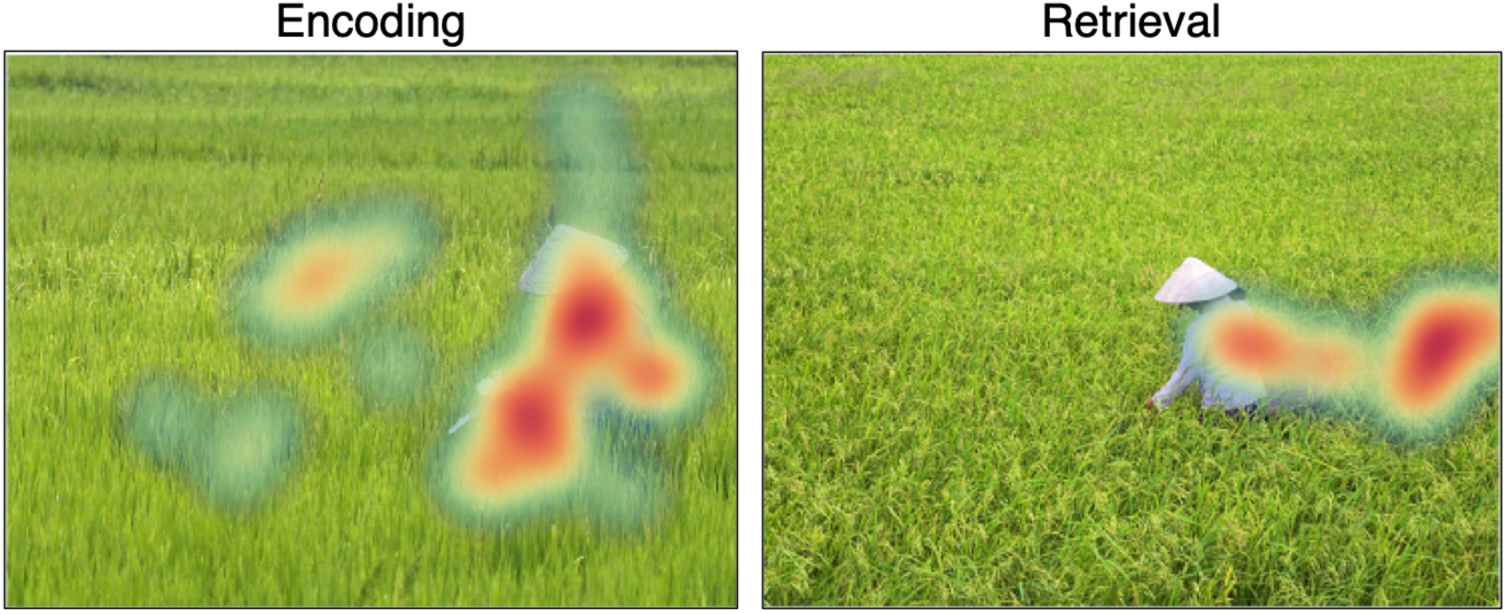
Visualization of EMs for a single participant during encoding and retrieval. In this example, the retrieval cue was a lure image presented at 40% degradation. The lure image is presented here for visualization purposes; it was not present on the screen during retrieval. Note that EMs at retrieval extend to the right of the central figure, to the location previously occupied by the figure in the encoded image.

**Fig 4.**
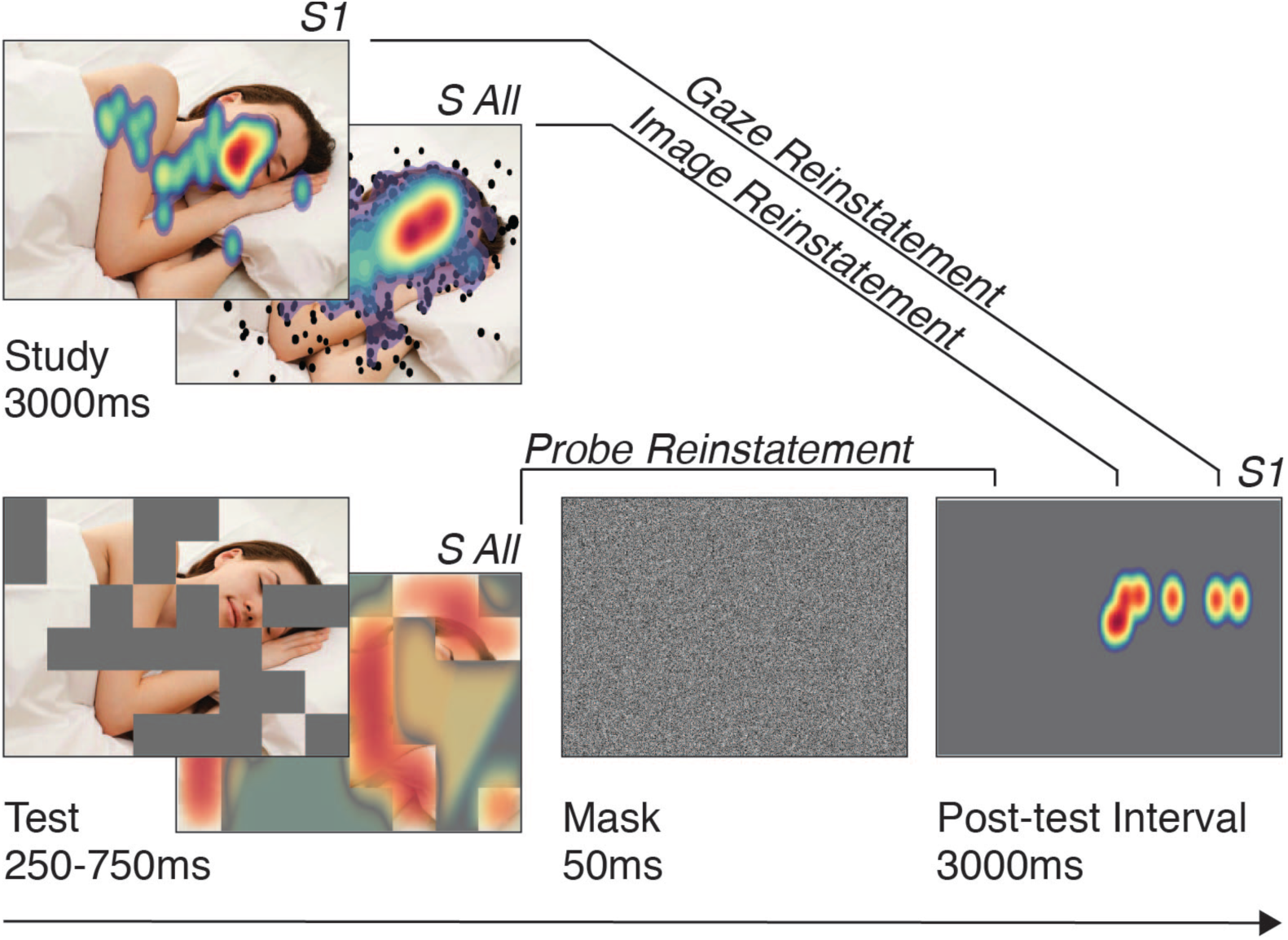
Illustration of the three measures of EM-based reinstatement. The heat maps reflect fixation density, with warm values indicating areas of high fixation density. Probe reinstatement is computed by correlating the heat map generated from the test probe weighted by the fixations of all participants (S All) viewing the same image during study and a single participant (S1) subsequently retrieving that image (or a similar lure) during the post-test interval. Image reinstatement is computed by correlating the heat map generated from the cumulative fixations of all participants (S All) viewing a single image over 4 study presentations and a single participant (S1) subsequently retrieving that image (or a similar lure) during the post-test interval. Gaze reinstatement is computed by correlating the heat map generated from the cumulative fixations of a single participant (S1) viewing a single image over 4 study presentations and subsequently retrieving that image (or a similar lure) during the post-test interval. All density maps are smoothed and duration-weighted.

To quantify the reinstatement of image features from LTM, we computed two additional measures of reinstatement. *Gaze reinstatement* reflects the similarity between the EM patterns of a single participant encoding a single image over four study opportunities and subsequently retrieving that same image (or a similar lure) during the 3s stimulus-free post-test interval. Based on the theory that EMs both encode and are themselves embedded in mnemonic representations (Noton & Stark, 1971a, 1971b), this measure captures reinstatement of the encoded image *including* the corresponding encoding operations, in this case, the pattern of EMs. Although there is ample research supporting a specialized role for such gaze reinstatement in memory retrieval (for review, see Wynn, Shen, et al., 2019), other work suggests that retrieval-related EMs reinstate salient regions of the encoded stimulus, even when EMs are restricted at encoding (Johansson et al., 2012). Therefore, to distinguish idiosyncratic reinstatement from more general image reinstatement, and to determine whether there is something special about reinstating one’s own fixations that benefits memory above and beyond the benefits conferred by reinstating generally salient or semantically informative regions of the encoded image, we also measured *image reinstatement*. This measure reflects the similarity between the EM patterns of *all* participants encoding a single image over the four study opportunities and a single participant subsequently retrieving that image during the 3s stimulus-free post-test interval, and captures reinstatement of the encoded image without requiring reinstatement of the accompanying EMs.

Fixations shorter than 80ms in duration, and fixations beginning before the start of the delay period, were removed from all analyses. Fixation density maps were duration-weighted and Gaussian-smoothed (σ = 80). Density maps were compared using a Pearson correlation, yielding a single value representing the spatial overlap between fixations during the post-test interval and fixations during study or test. To ensure that similarity values were driven by memory and not by participant-specific viewing patterns (e.g., scanning each image from left to right), we additionally correlated each post-test interval density map with 50 other randomly selected study (or test) image density maps. These resulting reinstatement scores were then averaged to obtain a control reinstatement score, which was subtracted from the reinstatement score of interest. In this way, all EM-based reinstatement measures control for image-invariant viewing tendencies, including the tendency to fixate the center of the screen. These measures are described in further detail below.

#### Gaze Reinstatement

To measure mnemonic reinstatement that is both participant- and image-specific, we generated density maps for each participant for each studied image using all fixations belonging to that participant viewing that image across all four blocks (four presentations of the same image; mean = 7.96 fixations per image presentation). The same method was applied to image retrieval, using all fixations made during the post-test interval (the 3s period between the presentation of the test probe and the response screen; mean = 5.36 fixations). For old images, the analysis proceeded as described, by comparing the gaze patterns of the same participant viewing and visualizing the same image during the study period and post-test interval, respectively (i.e., A-A). Importantly, for lure images presented at test, the post-test interval gaze pattern was compared to the gaze pattern of the same participant viewing the similar (lure) image during study (i.e., A-B). Thus, a high gaze reinstatement score for an old image indicates reinstatement of the same image, whereas a high score for a lure image indicates reinstatement of the similar studied image.

#### Image Reinstatement

While gaze reinstatement provides an index of the extent to which participants retrieve the encoded image by reproducing their own gaze pattern, it is possible that reinstatement during the post-test interval is participant-invariant. That is, retrieval-related gaze patterns might reflect retrieval of generally salient regions of the studied image, without reinstatement of the participant-specific associated EM pattern. Thus, in order to measure reinstatement of participant-invariant salient image regions, we generated duration-weighted smoothed density maps for each studied image using all fixations belonging to *all participants* viewing that image across all four blocks. These across-participant study density maps were compared to density maps reflecting the fixations made during the post-test interval by a single participant viewing a single image. By comparing gaze reinstatement to image reinstatement, we were able to distinguish between reinstatement that is image-specific from reinstatement that is participant- and image-specific and determine whether reinstating one’s own fixations benefits memory above and beyond reinstating generally salient regions of the remembered image.

#### Probe Reinstatement

Finally, in order to determine whether post-test interval reinstatement reflects information gleaned from the just-presented test probe, we computed the similarity of participant- and image-specific post-test interval fixations to a density map generated from the test probe image. Density maps were created by first generating pseudo-fixations for each pixel within the visible squares of the test probe and then creating a smoothed map (σ = 80) of those fixations (i.e., visible image squares) multiplied by the across-participant study density map for that image (see image reinstatement). This allowed for the visible squares of the test probe to be weighted by their respective saliency values, as determined by the gaze patterns of all participants who viewed that image during the study period. This procedure effectively downweights fixations, for example in visible squares on the periphery of the image (side, top, corners), that tend to attract few fixations. The resulting density map reflects the overall saliency of the visible portion of the test probe image.

### Data Analysis

To investigate factors contributing to performance on the recognition task, we ran a generalized linear mixed effects model (GLMM; glmer of package lme4, Bates, Mächler, Bolker, & Walker, 2015) with a bobyqa optimizer on trial-level accuracy (correct, incorrect, with a binomial distribution and logistic link function), and linear mixed effects models (LMEM) on gaze reinstatement, image reinstatement, and probe reinstatement, with probe type (old, lure), degradation (0%, 20%, 40%, 60%, 80%) and duration (250ms, 500ms, 750ms) as independent variables. To allow for simple effects analysis of significant interactions, probe type was recoded as 0 (old) and 1 (lure). Duration and degradation were z-scored and participant and item were modelled as random effects (intercepts). To build the models, we used a backward selection approach, starting with a maximal model (Bates et al., 2015) which included fixed effects for all variables and their interactions, as well as random intercepts for participant and item. Models were compared using likelihood ratio tests with α = 0.05, such that non-significant fixed effects were removed from the model in a stepwise fashion until no further model changes resulted in a significant likelihood ratio test. Results of the final best fit models arrived at via model comparison are reported, with significance values approximated with the lmerTest R package (Kuznetsova, Brockhoff, & Christensen, 2017).

To evaluate our hypotheses that gaze reinstatement should be positively predictive of accuracy for old images and negatively predictive of accuracy for lure images, we ran three additional GLMMs on accuracy by adding each of the described reinstatement scores to the final GLMM from the behavioral analysis. The models including reinstatement were compared to the base model using likelihood ratio tests with α = 0.05. Model comparison proceeded in a stepwise fashion, starting with main effects of reinstatement and proceeding to interactions of reinstatement and other predictors, such that only effects that significantly improved the fit of the model were retained.

To investigate individual differences in the relationship between gaze reinstatement and behavioral performance, we ran bootstrapped Pearson correlations (*bootstrap iterations* =5000) of gaze reinstatement and accuracy (overall % correct) separately for old and lure images. Finally, to examine changes in reinstatement over time, we ran three additional LMEMs for each 1000ms time bin within the post-test interval (0-1000ms, 1000-2000ms, 2000-3000ms) with density (i.e., reinstatement) value as the dependent variable and reinstatement measure (gaze reinstatement, image reinstatement, probe reinstatement), accuracy (correct, incorrect) and probe type (old, lure) as independent variables. Model parameters were the same as those reported above.

## Results

### Behavioral results

Results of the best fit model of accuracy arrived at via model comparison revealed significant effects of probe type, duration, and degradation (Table 1). As expected, memory accuracy was greater for old relative to lure images (Fig 5A), and increased with increased test probe duration (Fig 5B) and with decreased test probe degradation (Fig 5C).

**Fig 5.**
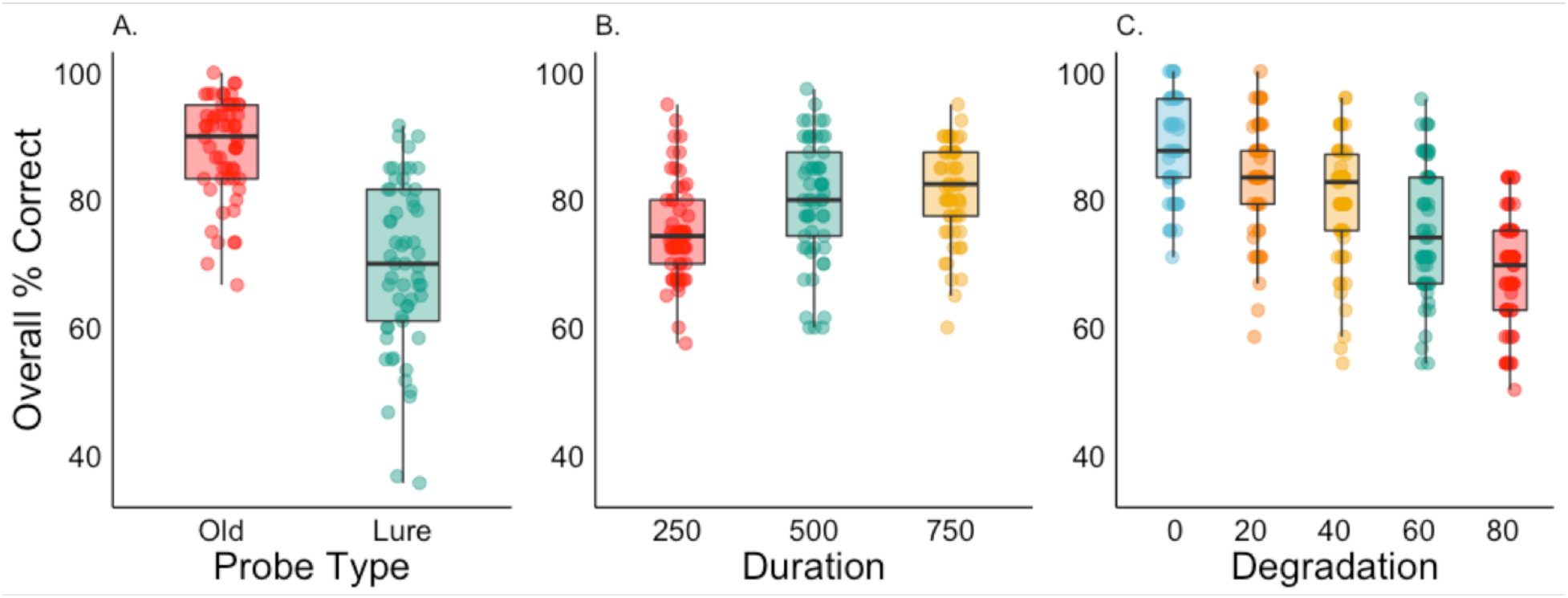
Accuracy (overall % correct) by (A) probe type, (B) duration, and (C) degradation.

**Table 1.**

| Table 1. |  |  |  |  |  |
| --- | --- | --- | --- | --- | --- |
| Fixed effects |  |  |  |  |  |
| | $\beta$ | SE | 95% CI | Z | p |
| Intercept | 2.317 | 0.085 | 2.153, 2.495 | 27.285 | <.001 *** |
| Duration | 0.184 | 0.032 | 0.119, 0.249 | 5.669 | <.001 *** |
| Degradation | - 0.472 | 0.044 | -0.561, -0.387 | -10.815 | <.001 *** |
| Probe Type | -1.342 | 0.069 | -1.485, -1.208 | -19.403 | <.001 *** |
| Total Observations = 6660 |  |  |  |  |  |
|  |  | Variance |  | SD |  |
| Random Effect for Participant (Intercept) |  | 0.156 |  | 0.394 |  |
| Random Effect for Item (Intercept) |  | 0.440 |  | 0.666 |  |
| Model equation: Accuracy ~ Duration + Degradation + Probe Type + (1 \| Participant) + (1 \| Image) |  |  |  |  |  |

### Eye movement results

To investigate the factors contributing to reinstatement of the test probe and the encoded image, we ran linear mixed effects models on probe reinstatement, image reinstatement, and gaze reinstatement, with duration, degradation, and probe type as predictors. We were particularly interested in the ability of each of these measures to differentiate between retrieval of old and lure images. For all analyses, reinstatement is reported as the difference between non-permuted and permuted similarity scores, such that positive scores indicate reinstatement of the same image that is greater than reinstatement of other images derived from a permutation.

#### Probe Reinstatement

Results of the best fit model of probe reinstatement (i.e., reinstatement of the test probe image, Table 4) arrived at via model comparison revealed significant effects of duration (Fig 6A) and degradation (Fig 6B), indicating that reinstatement of the test probe decreased with increased test probe duration and degradation.

**Fig 6.**
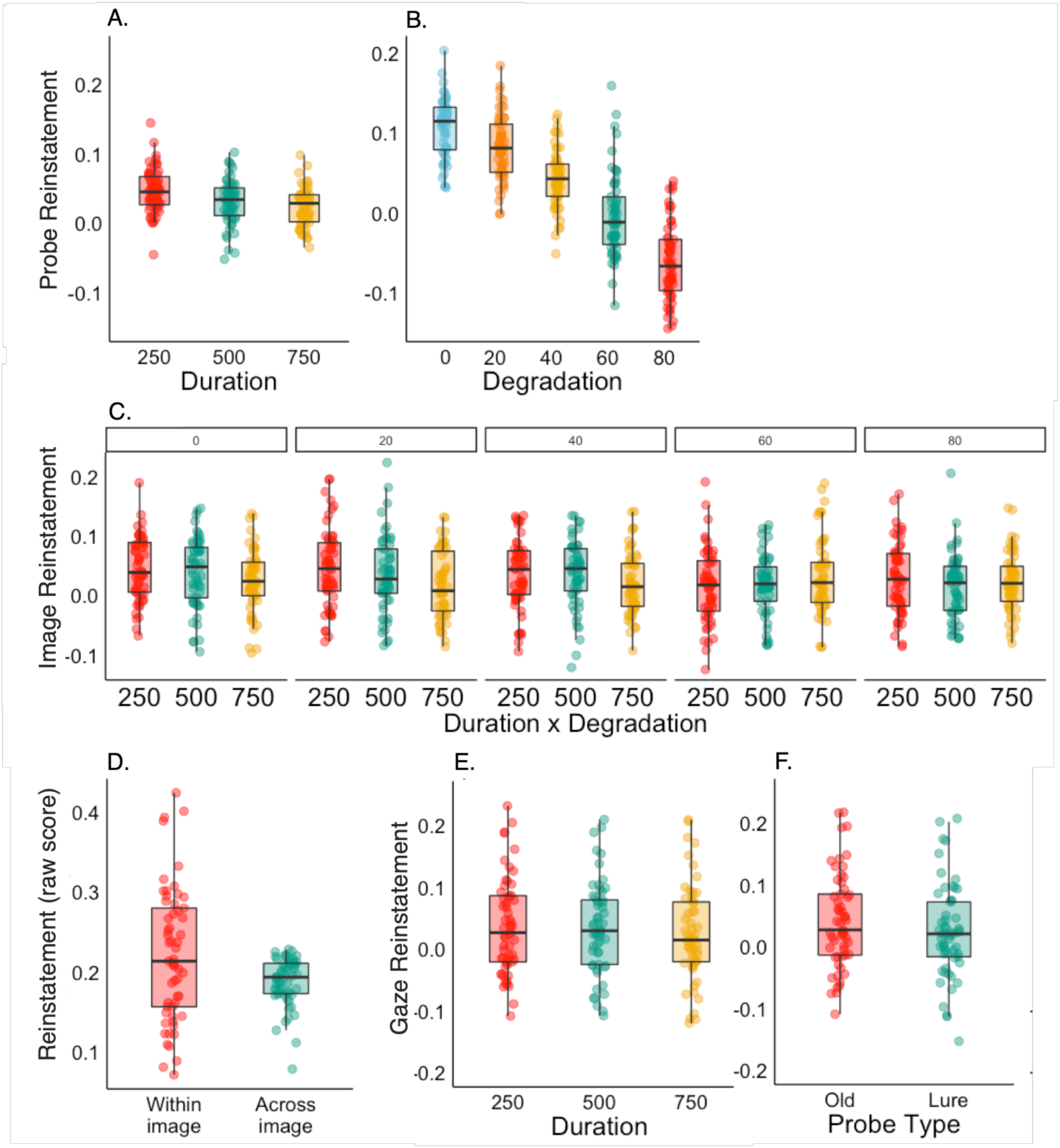
(A) Probe reinstatement by duration and (B) degradation. (C) Image reinstatement by duration and degradation. (D) Reinstatement scores within-image and across-image (permuted 50 times). Gaze reinstatement is computed by subtracting the across-image (permuted) score from the within-image score. (E) Gaze reinstatement by duration and (F) probe type.

#### Image Reinstatement

Results of the best fit model of image reinstatement (i.e., reinstatement of generally salient image regions, Table 3) arrived at via model comparison revealed a significant main effect of duration and a significant interaction of duration x degradation (Fig 6C), indicating that reinstatement decreased with increased test probe duration, and this effect was attenuated with increased test probe degradation.

#### Gaze Reinstatement

Before conducting the linear mixed effects model (LMEM) on gaze reinstatement, we first compared the reinstatement values (within-participant, within-image) to the permuted reinstatement values (within-participant, across-image) to ensure that retrieval-related gaze patterns were indeed closer to encoding-related gaze patterns from the same image than to encoding-related gaze patterns from different images. A paired samples *t*-test comparing reinstatement values (mean = 0.22) to permuted reinstatement values (mean = 0.19) was significant [*t* (56) = 3.4, *p* = .001, (Fig 6D)], indicating that reinstatement of encoding-related gaze patterns was indeed greater than would be expected based on idiosyncratic viewing tendencies.

Results of the best fit model of gaze reinstatement (i.e., reinstatement of own fixations, Table 2) arrived at via model comparison revealed a significant effect of duration (Fig 6E), indicating that reinstatement decreased with increased test probe duration. Notably, mean reinstatement was significantly greater than chance (0) for all levels of test probe duration (Fig 6E) and image degradation (*M_0_* = 0.030, *M_20_*= 0.039, *M_40_* = 0.037, *M_60_* = 0.025, *M_80_*= 0.034), suggesting that even briefly presented or severely degraded visual input is sufficient to elicit gaze reinstatement. Finally, whereas probe type was eliminated from both the probe reinstatement and image reinstatement models, we observed a significant effect of probe type on gaze reinstatement (Fig 6F), indicating that reinstatement of the encoded image *and* the corresponding encoding EMs differentiates retrieval of old images from retrieval of lure images.

**Table 2.**

| Table 2. |  |  |  |  |  |
| --- | --- | --- | --- | --- | --- |
| Fixed effects |  |  |  |  |  |
| | $\beta$ | SE | 95% CI | $t$ | $p$ |
| Intercept | 0.042 | 0.010 | 0.022, 0.062 | 4.144 | <.001 *** |
| Duration | -0.005 | 0.002 | -0.009, -0.000 | -2.081 | 0.038 * |
| Probe Type | -0.018 | 0.004 | -0.027, -0.010 | -4.223 | <.001 *** |
| Total Observations = 6803 |  |  |  |  |  |
|  |  | Variance |  | SD |  |
| Random Effect for Participant (Intercept) |  | 0.005 |  | 0.071 |  |
| Random Effect for Item (Intercept) |  | 0.005 |  | 0.068 |  |
| Model equation: Gaze reinstatement ~ Duration + Probe Type + (1 \| Participant) + (1 \| Image) |  |  |  |  |  |

**Table 3.**

| Table 3. |  |  |  |  |  |
| --- | --- | --- | --- | --- | --- |
| Fixed effects |  |  |  |  |  |
| | $\beta$ | SE | 95% CI | $T$ | $p$ |
| Intercept | 0.029 | 0.005 | 0.019, 0.040 | 5.09 | <.001 *** |
| Duration | -0.004 | 0.002 | -0.008, -0.001 | -2.702 | 0.007 ** |
| Degradation | -0.006 | 0.005 | -0.016, 0.004 | -1.124 | 0.262 |
| Duration x Degradation | 0.004 | 0.002 | 0.001, 0.008 | 2.598 | 0.009 ** |
| Total Observations = 6803 |  |  |  |  |  |
|  |  | Variance |  | SD |  |
| Random Effect for Participant (Intercept) |  | 0.000 |  | 0.009 |  |
| Random Effect for Item (Intercept) |  | 0.014 |  | 0.116 |  |
| Model equation: Image reinstatement ~ Duration x Degradation + (1 \| Participant) + (1 \| Image) |  |  |  |  |  |

**Table 4.**

| Table 4. |  |  |  |  |  |
| --- | --- | --- | --- | --- | --- |
| Fixed effects |  |  |  |  |  |
| | $\beta$ | SE | 95% CI | $T$ | $p$ |
| Intercept | 0.035 | 0.006 | 0.023, 0.046 | 5.923 | <.001 *** |
| Duration | -0.009 | 0.002 | -0.013, -0.005 | -4.864 | <.001 *** |
| Degradation | -0.060 | 0.006 | -0.071, -0.049 | -10.630 | <.001 *** |
| Total Observations = 6660 |  |  |  |  |  |
|  |  | Variance |  | SD |  |
| Random Effect for Participant (Intercept) |  | 0.000 |  | 0.012 |  |
| Random Effect for Item (Intercept) |  | 0.017 |  | 0.130 |  |
| Model equation: Probe reinstatement ~ Duration + Degradation + (1 \| Participant) + (1 \| Image) |  |  |  |  |  |

#### Summary of eye movement results

Results of the analyses on viewing behavior indicated that all three measures of reinstatement (probe reinstatement, image reinstatement, gaze reinstatement) decreased with increased test probe duration, suggesting that EMs may play a more significant role in retrieval when visual cues are insufficient. Degradation of the test probe significantly attenuated probe reinstatement, but had no effect on image or gaze reinstatement, further suggesting that sparse visual information is sufficient to elicit reinstatement of an encoded image from LTM, whereas reinstatement of the test probe is dependent on its visual properties. Finally, only gaze reinstatement showed a significant effect of probe type, with greater gaze reinstatement for old compared to lure test probes. To further investigate whether gaze reinstatement plays a specialized role in retrieval, we subsequently investigated the relationship between gaze reinstatement and memory performance in a model containing all reinstatement measures.

### Relationship between eye movements and behavior

To test our hypotheses that gaze reinstatement should be positively associated with accuracy for old images and negatively associated with accuracy for lure images, we investigated whether adding each of the previously described reinstatement scores to the accuracy model (Table 1) improved the model fit. As expected, the addition of gaze reinstatement significantly improved the fit of the model (*χ^2^* = 11.769, *p* < .001), whereas additions of image reinstatement (*χ^2^*= 2.921, *p* = .09) and probe reinstatement (*χ^2^* = 0.039, *p* = .843) did not. To account for encoding effects, we also added the cumulative number of fixations (z-scored) on each image during the study period (over the four study presentations) as a predictor. This measure has been previously linked to memory success (Chan, Kamino, Binns, & Ryan, 2011; Loftus & Mackworth, 1978; Olsen et al., 2016) and hippocampal activity (Liu et al., 2017). The addition of cumulative study fixations significantly improved the fit of the model (*χ^2^* = 10.354, *p* = .001). Interactions of gaze reinstatement and cumulative study fixations with the other predictors (probe type, duration, degradation), as well with each other, were subsequently added to the model in a stepwise manner. Only the interaction of gaze reinstatement and probe type significantly improved the fit of the model. The results of the best fit accuracy model (with the additions gaze reinstatement and cumulative study fixations) arrived at via model comparison are reported in Table 5.

**Table 5.**

| Table 5. |  |  |  |  |  |
| --- | --- | --- | --- | --- | --- |
| Fixed effects |  |  |  |  |  |
| | $\beta$ | SE | 95% CI | $z$ | $P$ |
| Intercept | 2.312 | 0.084 | 2.151, 2.489 | 27.499 | <.001 *** |
| Gaze Reinstatement | 0.050 | 0.059 | -0.068 0.169 | 0.841 | 0.401 |
| Probe Type | -1.346 | 0.069 | -1.489, -1.212 | -19.428 | <.001 *** |
| Duration | 0.179 | 0.032 | 0.114, 0.245 | 5.523 | <.001 *** |
| Degradation | -0.474 | 0.043 | -0.563, -0.390 | -10.923 | <.001 *** |
| Cumulative Study Fixations | 0.106 | 0.042 | 0.023, 0.189 | 2.557 | 0.011 * |
| Gaze Reinstatement x Probe Type | -0.242 | 0.071 | -0.385, -0.100 | -3.394 | <.001 *** |
| Total Observations = 6660 |  |  |  |  |  |
|  |  | Variance |  | SD |  |
| Random Effect for Participant (Intercept) |  | 0.148 |  | 0.385 |  |
| Random Effect for Item (Intercept) |  | 0.429 |  | 0.654 |  |
| Model Equation: Accuracy ~ Gaze Reinstatement + Probe Type + Duration + Degradation + Cumulative Study Fixations + Gaze Reinstatement x Probe Type + (1 \| Participant) + (1 \| Item) |  |  |  |  |  |

Results of the final best fit model of accuracy arrived at via model comparison (Table 5) revealed significant effects of probe type (old > lure), duration, and degradation, with accuracy increasing with increased test probe duration and decreased test probe degradation. In line with previous work (Chan et al., 2011; Loftus & Mackworth, 1978; Olsen et al., 2016), cumulative study fixations (a measure of encoding success) significantly predicted accuracy on the recognition task. Finally, the effect of gaze reinstatement on accuracy was non-significant for old images, and significantly negative for lure images, suggesting that reinstatement of participant- and image-specific encoding-related gaze patterns does not support recognition of old images, but does predict false alarms to similar (lure) images at test.

To determine whether the significant interaction of gaze reinstatement and probe type revealed by the GLMM was also present in an across-participants analysis, we ran bootstrapped Pearson correlations (*bootstrap iterations* =5000) of gaze reinstatement and accuracy (overall % correct) separately for old and lure images, see Fig 7A. As was the case in the within-participants analysis, we did not find evidence supporting the hypothesis that gaze reinstatement predicts correct recognition of old images (*r =* 0.053, 95% CI: −0.183, 0.291). However, in line with our predictions and results of the within-participants analysis, there was a robust negative correlation between gaze reinstatement and accuracy for lure images, suggesting that reinstatement of participant-and image-specific encoding-related gaze patterns during the post-test interval predicts false endorsement of similar images as old (*r =* −0.394, 95% CI: −0.660, −0.144).

**Fig 7.**
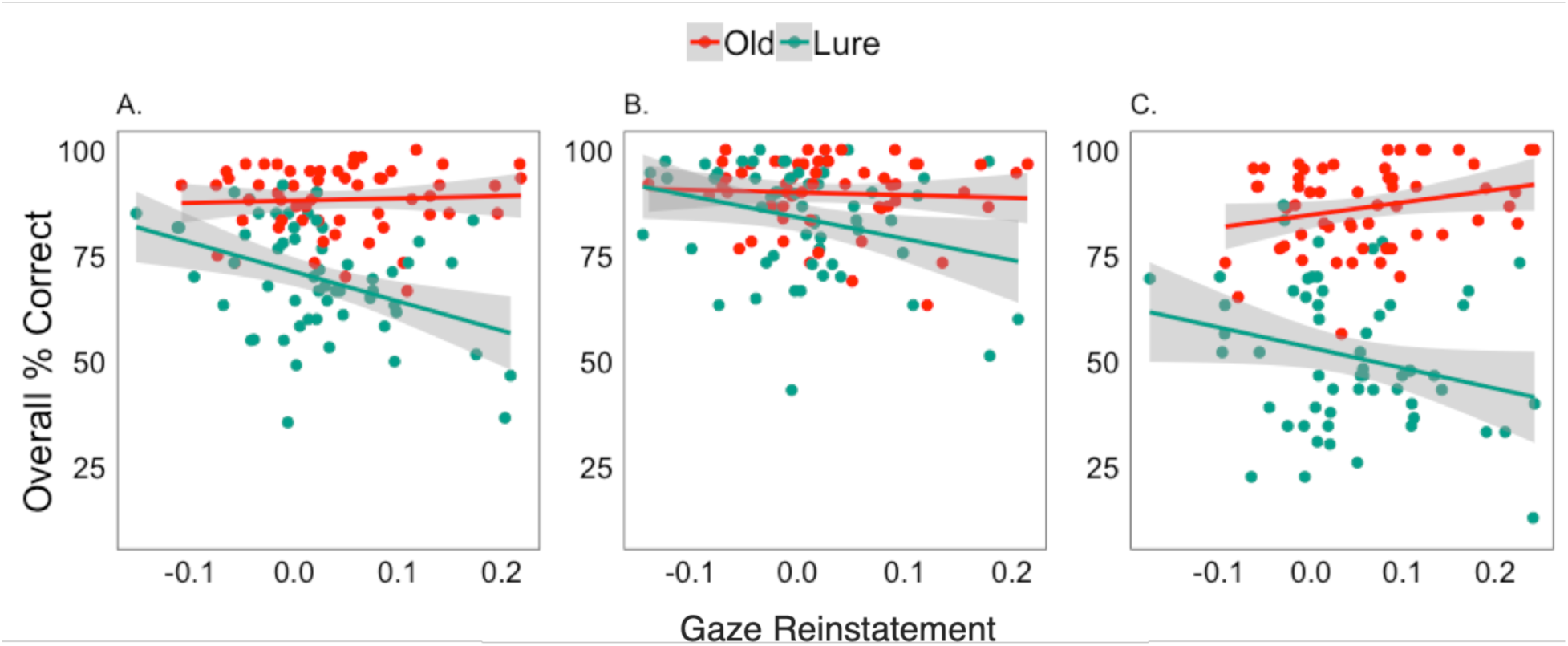
Correlations of old and lure gaze reinstatement and accuracy (overall % correct) for (A) all images, (B) easy images (median accuracy > 80.702), and (C) difficult images (median accuracy < 80.702).

Previous work suggests that gaze reinstatement might only facilitate memory performance when cognitive demands exceed cognitive resources (Wynn et al., 2018; for review, see Wynn, Shen, et al., 2019). Given that performance was at ceiling for old images, we were unable to probe this effect. Thus, to examine whether mnemonic demands modulated the effect of gaze reinstatement for old images, we conducted a median split of the data based on the mean accuracy for each image (median = 80.702), which we used to approximate difficulty in correctly recognizing the image (for related work on memorability, see Bylinskii, Isola, Bainbridge, Torralba, & Oliva, 2015). In line with the results of the previous bootstrap, gaze reinstatement was not significantly correlated with accuracy when only easy images were included in the analysis (*r = −*0.057, 95% CI: −0.304, 0.184), see Fig 7B. However, when only difficult images were included in the analysis, gaze reinstatement was positively correlated with accuracy for old images (*r =* 0.251, 95% CI: 0.029, 0.485), see Fig 7C.

### Reinstatement across time

To visualize the distinct trajectories of the three measures of reinstatement across the retrieval interval, we calculated density values for each measure (probe reinstatement, image reinstatement, gaze reinstatement) at discrete points in time from the onset of the test probe to the end of the post-test interval by sampling from each of the three density maps in 50ms intervals (collapsing across duration and degradation), see Fig 8A. This analysis differed slightly from the previous analysis in that rather than generating density maps for each time bin, we extracted the value of the respective density map (as described previously) at the location of fixation at each point in time (sampled in 50ms intervals).

**Fig 8.**
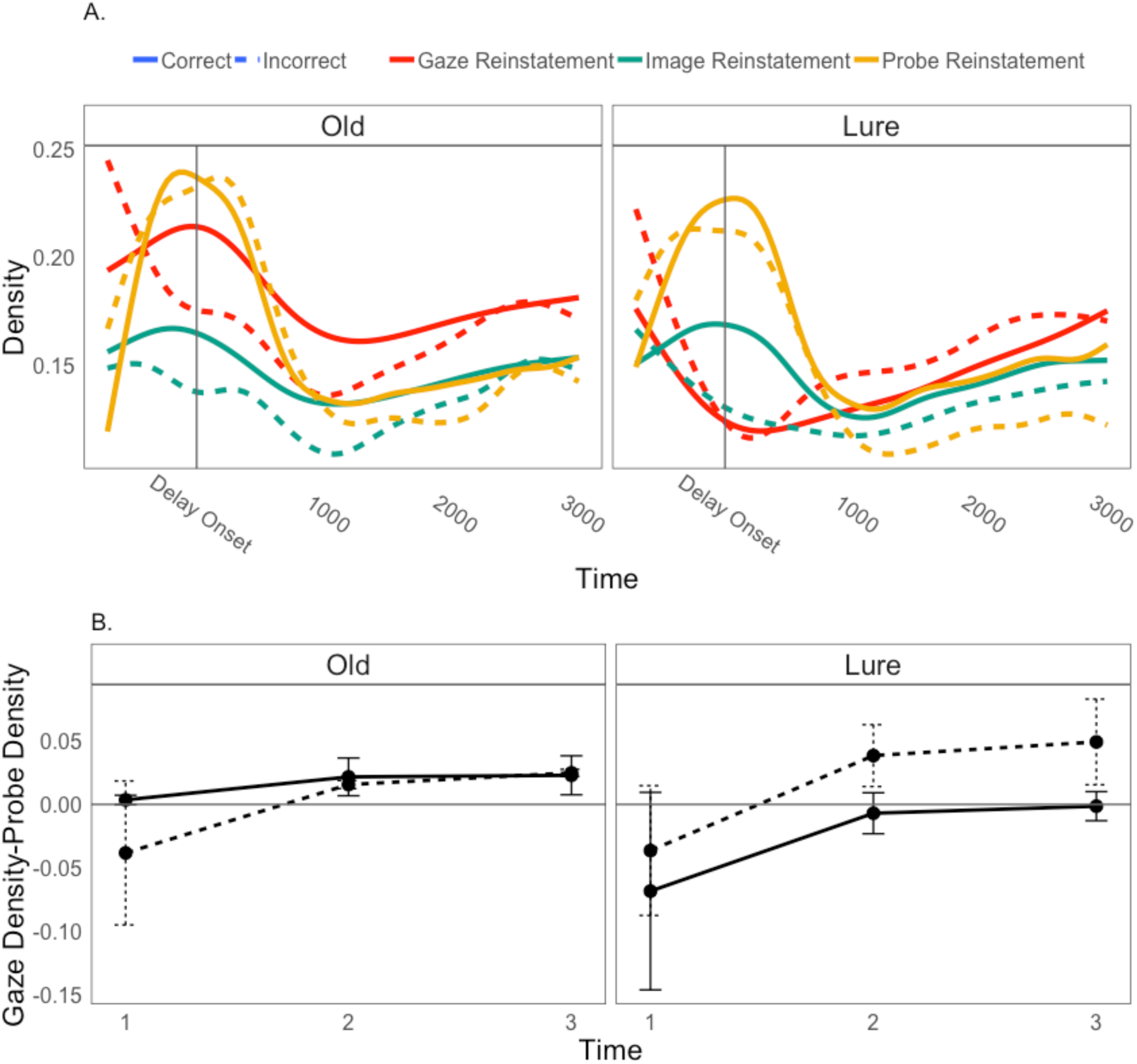
(A) Gaze reinstatement, image reinstatement, and probe reinstatement across time for old and lure images. Similarity is indexed by the value in the respective density map at each point (50ms) in time. Solid lines represent accurate responses and dotted lines represent inaccurate responses. (B) Mean difference scores (gaze density - probe density) for correct and incorrect old and lure images at each time bin. Error bars represent 95% CIs around the mean difference score.

For ease of analysis and interpretation when comparing density values across the post-test interval, we aggregated the values into three discrete 1000ms time bins. We ran separate LMEMs for old and lure images with density as the dependent variable and accuracy (correct, incorrect), measure (i.e., density map: gaze, image, probe), and time (T1: 0-1000ms, T2: 1000-2000ms, T3: 2000-3000ms) as independent variables. To allow for simple effects analysis of significant interactions, accuracy was coded as 0 (incorrect) and 1 (incorrect), time was coded for a linear effect, and gaze reinstatement was set the reference group for measure. Participant and item were modelled as random effects (intercepts). Models were compared using likelihood ratio tests with α = 0.05, such that non-significant fixed effects were removed from the model in a backward stepwise manner until no further model changes resulted in a significant likelihood ratio test. Results of the final best fit models arrived at via model comparison are reported below.

#### Old

Results of the final best fit model of density for old images arrived at via model comparison (Table 6) revealed a significant effect of accuracy for gaze reinstatement (reference category) at T1 (hits > misses), and this effect was marginally attenuated for image reinstatement and significantly attenuated for probe reinstatement. We also observed a significant interaction of accuracy and time, indicating that the difference in gaze reinstatement between hits and misses (hits > misses) significantly decreased over the course of the post-test interval. This effect (accuracy x time) was significantly attenuated and even reversed for probe reinstatement, indicating that over time, the difference in probe reinstatement between hits and misses (hits > misses) significantly increased relative to gaze reinstatement. Finally, probe reinstatement significantly outperformed gaze reinstatement for misses early in the post-test interval and this difference was significantly attenuated and even reversed over time (gaze > probe). For hits, the difference between probe reinstatement and gaze reinstatement was significantly attenuated at T1, however this difference significantly increased and was even reversed over time, such that gaze reinstatement outperformed probe reinstatement later in the post-test interval.

**Table 6.**

| Table 6. |  |  |  |  |  |
| --- | --- | --- | --- | --- | --- |
| Fixed Effects |  |  |  |  |  |
| | $\beta$ | SE | 95% CI | $t$ | $p$ |
| (Intercept) | 0.137 | 0.010 | 0.12, 0.16 | 14.06 | <.001 *** |
| Time | 0.006 | 0.005 | -0.00, 0.02 | 1.084 | 0.278 |
| Accuracy | 0.042 | 0.008 | 0.03, 0.06 | 5.592 | <.001 *** |
| Image Measure | -0.019 | 0.010 | -0.04, 0.00 | -1.911 | 0.056 |
| Probe Measure | 0.031 | 0.010 | 0.01, 0.05 | 3.147 | 0.002 ** |
| Time:Accuracy | -0.015 | 0.006 | -0.03, -0.00 | -2.637 | 0.008 ** |
| Time : Image Measure | 0.004 | 0.008 | -0.01, 0.02 | 0.570 | 0.569 |
| Time:Probe Measure | -0.032 | 0.008 | -0.05, -0.02 | -4.190 | <.001 *** |
| Accuracy:Image Measure | -0.018 | 0.010 | -0.04, 0.00 | -1.760 | 0.078 . |
| Accuracy:Probe Measure | -0.037 | 0.010 | -0.06, -0.02 | -3.572 | <.001 *** |
| Time:Accuracy:Image Measure | 0.006 | 0.008 | -0.01, 0.02 | 0.726 | 0.468 |
| Time:Accuracy:Probe Measure | 0.022 | 0.008 | 0.01, 0.04 | 2.725 | 0.006 ** |
| Total Observations = 30375 |  |  |  |  |  |
|  | Variance |  | SD |  |  |
| Random Effect for Participant (Intercept) | 0.002 |  | 0.044 |  |  |
| Random Effect for Item (Intercept) | 0.007 |  | 0.083 |  |  |
| Model equation: Density ~ Measure *Accuracy * Time + (1 \| Participant) + (1 \| Image) |  |  |  |  |  |

#### Lure

Results of the final best fit model of density for lure images arrived at via model comparison (Table 7) revealed a significant effect of accuracy for gaze reinstatement (reference category) at T1 (false alarms > correct rejections) and this effect was significantly attenuated for both image and probe reinstatement. Interactions of accuracy and time were eliminated from the model, indicating that the difference in reinstatement between correct rejections and false alarms remained consistent across the post-test interval. Gaze reinstatement significantly outperformed image reinstatement at T1 irrespective of accuracy, and this difference increased over time. However, gaze reinstatement performed significantly worse than probe reinstatement at T1. This difference (probe > gaze) was significantly attenuated and even reversed over time, such that gaze reinstatement later in the post-test interval outperformed probe reinstatement.

**Table 7.**

*Lure*
| Table 7. |  |  |  |  |  |
| --- | --- | --- | --- | --- | --- |
| Fixed Effects |  |  |  |  |  |
| | $\beta$ | SE | 95% CI | $t$ | $p$ |
| (Intercept) | 0.145 | 0.007 | 0.13, 0.16 | 21.99 | <.001 *** |
| Time | 0.017 | 0.002 | 0.01, 0.02 | 9.548 | <.001 *** |
| Accuracy | -0.035 | 0.004 | -0.04, -0.03 | -9.678 | <.001 *** |
| Image Measure | -0.011 | 0.005 | -0.02, -0.00 | -2.481 | 0.031 * |
| Probe Measure | 0.019 | 0.005 | 0.01, 0.03 | 4.115 | <.001 *** |
| Time:Image Measure | -0.017 | 0.003 | -0.02, -0.01 | -6.419 | <.001 *** |
| Time:Probe Measure | -0.036 | 0.003 | -0.04, -0.03 | -14.04 | <.001 *** |
| Accuracy:Image Measure | 0.041 | 0.005 | 0.03, 0.05 | 8.952 | <.001 *** |
| Accuracy:Probe Measure | 0.043 | 0.005 | 0.03, 0.05 | 9.365 | <.001 *** |
| Total Observations = 30462 |  |  |  |  |  |
|  | Variance |  | SD |  |  |
| Random Effect for Participant (Intercept) | 0.001 |  | 0.037 |  |  |
| Random Effect for Item (Intercept) | 0.004 |  | 0.067 |  |  |
| Model equation: Density ~ Measure * Time + Measure * Accuracy + (1 \| Participant) + (1 \| Image) |  |  |  |  |  |

#### Summary of temporal analyses

In summary, results of the temporal analyses revealed changes in the spatial distribution of EMs over time as well as distinct gaze trajectories for old and lure images over the post-test interval. For both old and lure images, early fixations returned to regions of the screen previously occupied by salient regions of the test probe. However, with increasing time, EMs were primarily directed to regions that were previously occupied by salient regions of the studied image, and particularly those that were fixated previously (i.e., gaze reinstatement). These findings suggest that whereas initial EMs during retrieval may be biased by sensory input held online, subsequent EMs are increasingly driven by mnemonic processes. Moreover, gaze reinstatement was related to the subsequent memory judgment, with greater gaze reinstatement for hits compared to misses early in the post-test interval and for false alarms compared to correct rejections across the post-test interval. Together, these findings suggest that EMs flexibly support memory retrieval by differentially reinstating varying levels of mnemonic content across time and space.

## Discussion

Pattern completion describes a neurocomputational process whereby a memory representation is retrieved from partial or degraded input, and is a critical function of the hippocampal relational memory system (Marr, 1971; McClelland et al., 1995; for review, see Hunsaker & Kesner, 2013). Yet, it is often studied behaviorally. In mnemonic discrimination tasks, participants are required to discriminate repeated old items from similar “lure” items, with false endorsement of lure items taken as *prima facie* evidence of pattern completion (Bakker et al., 2008; Clelland et al., 2009; Stark et al., 2013; Toner et al., 2009). However, the relationship between this behavioral response and the underlying retrieval mechanism remains unclear. In the present study, we used an EM-based measure of encoding-retrieval overlap to investigate whether incomplete lure memory cues elicit retrieval of the originally encoded stimulus (i.e., that pattern completion has occurred). Based on evidence of behavioral pattern completion and EM-based reinstatement, we predicted that the similarity between encoding- and retrieval-related EMs or, *gaze reinstatement*, a measure that has been previously linked to memory retrieval (for review, see Wynn, Shen, et al., 2019), hippocampal activity (Ryals et al., 2015), and neural reactivation (Bone et al., 2018), would be spontaneously recruited in response to degraded test probes, and would be greater for hits than misses and for lure false alarms compared with correct rejections.

Consistent with previous findings of gaze reinstatement (Bone et al., 2018; Foulsham & Kingstone, 2013; Holm & Mäntylä, 2007; Johansson & Johansson, 2013; Kumcu & Thompson, 2018; Laeng et al., 2014; Scholz et al., 2016; for review, see Wynn, Shen, et al., 2019), retrieval-related EMs were more similar to the EMs enacted during encoding of the same (old) or similar (lure) image than to the EMs enacted during encoding of other images, suggesting that they reflect image-specific memory. In line with our predictions, gaze reinstatement was significantly greater than chance at all levels of test probe degradation, indicating that given an incomplete cue, EMs facilitate reactivation of a specific item representation from memory. Moreover, temporal analyses indicated that whereas early fixations following the test probe largely reinstated salient regions of the test probe, later fixations favored regions previously visited during encoding of the same or similar image. To investigate whether this reinstatement played a functional role in retrieval, we modeled accuracy on the recognition memory task as a function of EMs during both encoding and retrieval.

Consistent with findings of behavioral pattern completion (Bakker et al., 2008; Clelland et al., 2009; Stark et al., 2013; Toner et al., 2009), gaze reinstatement for lure images, that is, the similarity between EMs following presentation of a lure test probe and EMs during encoding of a similar image, was negatively correlated with accuracy both within and across participants. This effect could not be accounted for by reinstatement of generally salient regions of the studied image (i.e., image reinstatement), reinstatement of the presented test probe (i.e., probe reinstatement), or encoding-related EMs. Extending previous work, these findings lend critical support to the assumption that false alarms to lure items (i.e., behavioral pattern completion) reflect retrieval of previously encoded similar items (Bakker et al., 2008; Clelland et al., 2009; Stark et al., 2013; Toner et al., 2009). Moreover, these findings are consistent with, and may be related to, a growing literature suggesting that reinstatement of encoding-related neural activity patterns can have detrimental effects on recognition memory, particularly with respect to discriminating lures (e.g., Chadwick et al., 2016; Staudigl, Vollmar, Noachtar, & Hanslmayr, 2015; Ye et al., 2016).

In addition to the hypothesized role of gaze reinstatement in lure false alarms, we predicted that gaze reinstatement would facilitate correct recognition of old images, consistent with theories and findings of EM-based mnemonic reinstatement (Noton & Stark, 1971a, 1971b; for review, see Wynn, Shen, et al., 2019). Indeed, of the three reinstatement measures, only gaze reinstatement was significantly different between old images and lure images, suggesting that reinstatement of one’s own encoding-related EMs reflects the contents of memory. Whereas we did not observe a general effect of gaze reinstatement on recognition of old images, finer-grained temporal analyses revealed that within participants, the relationship between gaze reinstatement and performance emerged early in the post-test interval and was attenuated over time. Consistent with previous work, this finding suggests that recognition is more likely when early fixations reinstate remembered image regions (Foulsham & Kingstone, 2013; Holm & Mäntylä, 2007; Wynn et al., 2016). Moreover, analyses of individual differences revealed a positive correlation between gaze reinstatement and performance for difficult old images. This finding lends support to previous work suggesting that gaze reinstatement facilitates performance when task demands exceed cognitive resources (Wynn et al., 2018). Finally, results of the accuracy model further revealed a significant positive relationship between the number of cumulative fixations made across the four study presentations of each image and accuracy on the recognition task. This finding is consistent with previous work linking encoding-related EMs to memory performance (Chan et al., 2011; Loftus & Mackworth, 1978; Olsen et al., 2016) and hippocampal activity (Liu et al., 2017), and further suggests that EMs during both encoding and retrieval play a critical role in memory success.

Although the present study did not measure neural activity, several lines of evidence support the idea that the observed EM effect is related to hippocampal pattern completion. First, gaze reinstatement is increased in older adults, a population with deficits in hippocampal structure and function (for review, see Grady & Ryan, 2017), compared with younger adults (Wynn et al., 2016, 2018), and this is consistent with an age-related bias towards pattern completion (e.g., lure false alarms, Stark, Stevenson, Wu, Rutledge, & Stark, 2015). Indeed, a recent study from Vieweg et al. (2015) showed that the similarity between gaze patterns for correctly identified degraded images (e.g., library called library) and response matching incorrectly identified degraded images (e.g., office called library) was greater in older adults compared to younger adults and was correlated with an age-related bias towards behavioral pattern completion (learned images > new images). Second, lure false alarms are increased in cases of hippocampal damage due to mild cognitive impairment (Stark et al., 2013) or amnesia (Baker et al., 2016; Brock Kirwan et al., 2012). Finally, both gaze reinstatement (Ryals et al., 2015) and other retrieval-related EM effects (Bridge, Cohen, & Voss, 2017; Hannula & Ranganath, 2009; Ryan, Althoff, Whitlow, & Cohen, 2000) have been linked to activity in the hippocampus. Considered together with the present results, these findings suggest that the outcome of behavioral pattern completion (i.e., lure false alarms) might be attributed, at least in part, to the hippocampally-mediated reactivation of relational information via gaze reinstatement.

In summary, the present study used EM monitoring to provide evidence that lure false alarms are the result of pattern completion, whereby, in response to the presentation of an incomplete test probe, a visually similar, previously encoded image is retrieved. Given a partial retrieval cue, participants reinstated their encoding-related EMs for the same (old) or similar (lure) image, and this gaze reinstatement both supported the recognition of old images and increased the likelihood of false alarms for lure images. Over the course of retrieval, gaze trajectories shifted from initially reinstating just-viewed sensory input to subsequently reinstating image features and the accompanying EMs from memory, demonstrating that the component processes involved in EM-based reinstatement may flexibly adapt and change over time. Critically, these findings extend previous work by showing first, that gaze reinstatement influences subsequent behavioral pattern completion (i.e., lure false alarms), and second, that the “pattern” of input that is completed or reactivated during retrieval involves not only the stimulus itself (i.e., image reinstatement), but also the operation by which that stimulus was encoded (i.e., gaze reinstatement). Thus, taken together, the results of the present study suggest that gaze reinstatement supports behavioral pattern completion as part of its larger role in memory retrieval. More specifically, by re-enacting an encoded sequence of EMs, gaze reinstatement facilitates the reactivation of encoded stimulus features and the relations among them, which together constitute a “pattern” that can be subsequently “completed” or retrieved given a partial input cue. Further research should continue to explore the various ways in which EMs might support retrieval of complex stimuli and how information retrieved via gaze reinstatement might guide adaptive behavior.

## Acknowledgements

We would like to thank Jonathan Musat, Fahad Ahmad, Sam Alain and Rahim Ahmed for their assistance with recruitment and testing, as well as Tarek Amer for his helpful comments on earlier versions of the manuscript.

## Footnotes

1 This participant also had an average performance score lower than 1.5SD from the mean.

2 These participants also had average performance scores lower than 2SD from the mean.

## References

Armson, M. J., Ryan, J. D., & Levine, B. (2019). Maintaining fixation does not increase demands on working memory relative to free viewing. PeerJ, 7, e6839. https://doi.org/10.7717/peerj.6839

Baker, S., Vieweg, P., Gao, F., Gilboa, A., Wolbers, T., Black, S. E., & Rosenbaum, R. S. (2016). The Human Dentate Gyrus Plays a Necessary Role in Discriminating New Memories. Current Biology, 26(19), 2629–2634. https://doi.org/10.1016/j.cub.2016.07.081

Bakker, A., Kirwan, C. B., Miller, M., & Stark, C. E. L. (2008). Pattern separation in the human hippocampal CA3 and dentate gyrus. Science (New York, N.Y.), 319(5870), 1640–1642. https://doi.org/10.1126/science.1152882

Bates, D., Mächler, M., Bolker, B., & Walker, S. (2015). Fitting Linear Mixed-Effects Models Using lme4. Journal of Statistical Software, 67(1). https://doi.org/10.18637/jss.v067.i01

Bicanski, A., & Burgess, N. (2019). A Computational Model of Visual Recognition Memory via Grid Cells. Current Biology, 1–12. https://doi.org/10.1016/j.cub.2019.01.077

Bochynska, A., & Laeng, B. (2015). Tracking down the path of memory: eye scanpaths facilitate retrieval of visuospatial information. Cognitive Processing, 16(1), 159–163. https://doi.org/10.1007/s10339-015-0690-0

Bone, M. B., St-Laurent, M., Dang, C., McQuiggan, D. A., Ryan, J. D., & Buchsbaum, B. R. (2018). Eye Movement Reinstatement and Neural Reactivation During Mental Imagery. Cerebral Cortex, (July), 1–15. https://doi.org/10.1093/cercor/bhy014

Bridge, D. J., Cohen, N. J., & Voss, J. L. (2017). Distinct hippocampal versus frontoparietal-network contributions to retrieval and memory-guided exploration Donna. Journal of Cognitive Neuroscience, 26(3), 194–198. https://doi.org/10.1162/jocn

Brock Kirwan, C., Hartshorn, A., Stark, S. M., Goodrich-Hunsaker, N. J., Hopkins, R. O., & Stark, C. E. L. (2012). Pattern separation deficits following damage to the hippocampus. Neuropsychologia, 50(10), 2408–2414. https://doi.org/10.1016/j.neuropsychologia.2012.06.011

Bylinskii, Z., Isola, P., Bainbridge, C., Torralba, A., & Oliva, A. (2015). Intrinsic and extrinsic effects on image memorability. Vision Research, 116, 165–178. https://doi.org/10.1016/j.visres.2015.03.005

Chadwick, M. J., Anjum, R. S., Kumaran, D., Schacter, D. L., Spiers, H. J., & Hassabis, D. (2016). Semantic representations in the temporal pole predict false memories. Proceedings of the National Academy of Sciences, 113(36), 10180–10185. https://doi.org/10.1073/pnas.1610686113

Chan, J. P. K., Kamino, D., Binns, M. a, & Ryan, J. D. (2011). Can changes in eye movement scanning alter the age-related deficit in recognition memory? Frontiers in Psychology, 2(May), 1–11. https://doi.org/10.3389/fpsyg.2011.00092

Clelland, C. D., Choi, M., Romberg, C., Clemenson, G. D., Fragniere, A., Tyers, P., … Bussey, T. J. (2009). A Functional Role for Adult Hippocampal Neurogenesis in Spatial Pattern Separation. Science, 325(5937), 210–213. https://doi.org/10.1126/science.1173215

Cohen, N. J., & Eichenbaum, H. (1993). Memory, amnesia, and the hippocampal system. Memory, amnesia, and the hippocampal system. Cambridge, MA, US: The MIT Press.

Cowell, R. A., Barense, M. D., & Sadil, P. S. (2019). A roadmap for understanding memory: Decomposing cognitive processes into operations and representations. Eneuro, ENEURO.0122-19.2019. https://doi.org/10.1523/eneuro.0122-19.2019

Damiano, C., & Walther, D. B. (2019). Distinct roles of eye movements during memory encoding and retrieval. Cognition, 184(December 2018), 119–129. https://doi.org/10.1016/j.cognition.2018.12.014

Eichenbaum, H., & Cohen, N. J. (2001). From conditioning to conscious recollection: Memory systems of the brain. From conditioning to conscious recollection: Memory systems of the brain. New York, NY, US: Oxford University Press.

Ferreira, F., Apel, J., & Henderson, J. M. (2008). Taking a new look at looking at nothing. Trends in Cognitive Sciences, 12(11), 405–410. https://doi.org/10.1016/j.tics.2008.07.007

Foulsham, T., & Kingstone, A. (2013). Where have eye been? Observers can recognise their own fixations. Perception, 42(10), 1085–1089. https://doi.org/10.1068/p7562

Grady, C. L., & Ryan, J. D. (2017). Age-Related Differences in the Human Hippocampus: Behavioral, Structural and Functional Measures. In D. E. Hannula & M. C. Duff (Eds.), The Hippocampus from Cells to Systems (pp. 167–208). Cham: Springer International Publishing. https://doi.org/10.1007/978-3-319-50406-3_7

Hannula, D. E., Althoff, R. R., Warren, D. E., Riggs, L., & Cohen, N. J. (2010). Worth a glance: using eye movements to investigate the cognitive neuroscience of memory. Frontiers in Human Neuroscience, 4(October), 1–16. https://doi.org/10.3389/fnhum.2010.00166

Hannula, D. E., & Ranganath, C. (2009). The eyes have it: hippocampal activity predicts expression of memory in eye movements. Neuron, 63(5), 592–599. https://doi.org/10.1016/j.neuron.2009.08.025

Hannula, D. E., Ryan, J. D., Tranel, D., & Cohen, N. J. (2007). Rapid onset relational memory effects are evident in eye movement behavior, but not in hippocampal amnesia. Journal of Cognitive Neuroscience, 19(10), 1690–1705. https://doi.org/10.1162/jocn.2007.19.10.1690

Hannula, D. E., Ryan, J. D., & Warren, D. E. (2017). Beyond Long-Term Declarative Memory: Evaluating Hippocampal Contributions to Unconscious Memory Expression, Perception, and Short-Term Retention. In The Hippocampus from Cells to Systems (pp. 281–336). Cham: Springer International Publishing. https://doi.org/10.1007/978-3-319-50406-3_10

Henderson, J. M., Williams, C. C., & Falk, R. J. (2005). Eye movements are functional during face learning. Memory & Cognition, 33(1), 98–106. https://doi.org/10.3758/BF03195300

Holm, L., & Mäntylä, T. (2007). Memory for scenes: refixations reflect retrieval. Memory & Cognition, 35(7), 1664–1674. https://doi.org/10.3758/BF03193500

Hunsaker, M. R., & Kesner, R. P. (2013). The operation of pattern separation and pattern completion processes associated with different attributes or domains of memory. Neuroscience and Biobehavioral Reviews, 37(1), 36–58. https://doi.org/10.1016/j.neubiorev.2012.09.014

Johansson, R., Holsanova, J., Dewhurst, R., & Holmqvist, K. (2012). Eye movements during scene recollection have a functional role, but they are not reinstatements of those produced during encoding. Journal of Experimental Psychology. Human Perception and Performance, 38(5), 1289–1314. https://doi.org/10.1037/a0026585

Johansson, R., & Johansson, M. (2013). Look here, eye movements play a functional role in memory retrieval. Psychological Science, 25(1), 236–242. https://doi.org/10.1177/0956797613498260

Kragel, J. E., Vanhaerents, S., Templer, J. W., Schuele, S., Joshua, M., Nilakantan, A. S., & Bridge, D. J. (2019). Hippocampal theta coordinates memory processing during visual exploration.

Kumcu, A., & Thompson, R. L. (2018). Less imageable words lead to more looks to blank locations during memory retrieval. Psychological Research, 0(0), 0. https://doi.org/10.1007/s00426-018-1084-6

Kuznetsova, A., Brockhoff, P. B., & Christensen, R. H. B. (2017). lmerTest Package: Tests in Linear Mixed Effects Models. Journal of Statistical Software, 82(13). https://doi.org/10.18637/jss.v082.i13

Laeng, B., Bloem, I. M., D’Ascenzo, S., & Tommasi, L. (2014). Scrutinizing visual images: The role of gaze in mental imagery and memory. Cognition, 131(2), 263–283. https://doi.org/10.1016/j.cognition.2014.01.003

Laeng, B., & Teodorescu, D.-S. (2002). Eye scanpaths during visual imagery reenact those of perception of the same visual scene. Cognitive Science, 26(2), 207–231. https://doi.org/10.1207/s15516709cog2602_3

Liu, Z., Shen, K., Olsen, R. K., & Ryan, J. D. (2017). Visual sampling predicts hippocampal activity. Journal of Neuroscience, 37, 1–11. https://doi.org/10.1523/JNEUROSCI.2610-16.2016

Loftus, G. R., & Mackworth, N. H. (1978). Cognitive determinants of fixat ion locati on during picture viewing. Journal of Experimental Psychology: Human Perception and Performance, 4(4), 565 – 572.

Marr, D. (1971). Simple Memory: A Theory for Archicortex. Philosophical Transactions of the Royal Society B: Biological Sciences, 262(841), 23–81. https://doi.org/10.1098/rstb.1971.0078

McClelland, J., O’Reilly, R. C., & McNaughton, B. L. (1995). Why there are complimentary learning systems in the hippocampus and neocortex: insights from the successes and failures of connectionist models of learning and memory.

Molitor, R. J., Ko, P. C., Hussey, E. P., & Ally, B. a. (2014). Memory-related eye movements challenge behavioral measures of pattern completion and pattern separation. Hippocampus, 24(6), 666–672. https://doi.org/10.1002/hipo.22256

Nau, M., Julian, J. B., & Doeller, C. F. (2018). How the Brain’s Navigation System Shapes Our Visual Experience. Trends in Cognitive Sciences, xx, 1–16. https://doi.org/10.1016/j.tics.2018.06.008

Noton, D., & Stark, L. (1971a). Scanpaths in eye movements during pattern perception. Science (New York, N.Y.), 171(3968), 308–311. Retrieved from http://www.ncbi.nlm.nih.gov/pubmed/5538847

Noton, D., & Stark, L. (1971b). Scanpaths in saccadic eye movements while viewing and recognizing patterns. Vision Research, 11(9), 929–IN8. https://doi.org/10.1016/0042-6989(71)90213-6

Olsen, R. K., Chiew, M., Buchsbaum, B. R., & Ryan, J. D. (2014). The relationship between delay period eye movements and visuospatial memory. Journal of Vision, 14(1), 8–8. https://doi.org/10.1167/14.1.8

Olsen, R. K., Sebanayagam, V., Lee, Y., Moscovitch, M., Grady, C. L., Rosenbaum, R. S., & Ryan, J. D. (2016). The relationship between eye movements and subsequent recognition: Evidence from individual differences and amnesia. Cortex, 85, 182–193. https://doi.org/10.1016/j.cortex.2016.10.007

Rolls, E. T. (2013). The mechanisms for pattern completion and pattern separation in the hippocampus. Frontiers in Systems Neuroscience, 7(October), 1–21. https://doi.org/10.3389/fnsys.2013.00074

Ryals, A. J., Wang, J. X., Polnaszek, K. L., & Voss, J. L. (2015). Hippocampal contribution to implicit configuration memory expressed via eye movements during scene exploration. Hippocampus, 25(9), 1028–1041. https://doi.org/10.1002/hipo.22425

Ryan, J. D., Althoff, R. R., Whitlow, S., & Cohen, N. J. (2000). Amnesia is a Deficit in Relational Memory. Psychological Science, 11(6), 454–461. https://doi.org/10.1111/1467-9280.00288

Ryan, J. D., Shen, K., Kacollja, A., Tian, H., Griffiths, J., & McIntosh, R. (2018). The functional reach of the hippocampal memory system to the oculomotor system. BioRxiv. https://doi.org/https://doi.org/10.1101/303511

Santoro, A. (2013). Reassessing pattern separation in the dentate gyrus. Frontiers in Behavioral Neuroscience, 7(July), 1–4. https://doi.org/10.3389/fnbeh.2013.00096

Scholz, A., Mehlhorn, K., & Krems, J. F. (2016). Listen up, eye movements play a role in verbal memory retrieval. Psychological Research, 80(1), 149–158. https://doi.org/10.1007/s00426-014-0639-4

Shen, K., Bezgin, G., Selvam, R., McIntosh, A. R., & Ryan, J. D. (2016). An Anatomical Interface between Memory and Oculomotor Systems. Journal of Cognitive Neuroscience, 28(11), 1772–1783. https://doi.org/10.1162/jocn_a_01007

Stark, S. M., Stevenson, R., Wu, C., Rutledge, S., & Stark, C. E. L. (2015). Stability of age-related deficits in the mnemonic similarity task across task variations. Behavioral Neuroscience, 129(3), 257–268. https://doi.org/10.1037/bne0000055

Stark, S. M., Yassa, M. A., Lacy, J. W., & Stark, C. E. L. (2013). A task to assess behavioral pattern separation (BPS) in humans: Data from healthy aging and mild cognitive impairment. Neuropsychologia, 51(12), 2442–2449. https://doi.org/10.1016/j.neuropsychologia.2012.12.014

Staudigl, T., Vollmar, C., Noachtar, S., & Hanslmayr, S. (2015). Temporal-Pattern Similarity Analysis Reveals the Beneficial and Detrimental Effects of Context Reinstatement on Human Memory. Journal of Neuroscience, 35(13), 5373–5384. https://doi.org/10.1523/JNEUROSCI.4198-14.2015

Toner, C. K., Pirogovsky, E., Kirwan, C. B., & Gilbert, P. E. (2009). Visual object pattern separation deficits in nondemented older adults. Learning & Memory (Cold Spring Harbor, N.Y.), 16(5), 338–342. https://doi.org/10.1101/lm.1315109

Vieweg, P., Stangl, M., Howard, L. R., & Wolbers, T. (2015). Changes in pattern completion – A key mechanism to explain age-related recognition memory deficits? Cortex, 64(September), 343–351. https://doi.org/10.1016/j.cortex.2014.12.007

Voss, J. L., Bridge, D. J., Cohen, N. J., & Walker, J. A. (2017). A Closer Look at the Hippocampus and Memory. Trends in Cognitive Sciences, 21(8), 577–588. https://doi.org/10.1016/j.tics.2017.05.008

Wynn, J. S., Bone, M. B., Dragan, M. C., Hoffman, K. L., Buchsbaum, B. R., & Ryan, J. D. (2016). Selective scanpath repetition during memory-guided visual search. Visual Cognition, 24(1), 15–37. https://doi.org/10.1080/13506285.2016.1175531

Wynn, J. S., Olsen, R. K., Binns, M. A., Buchsbaum, B. R., & Ryan, J. D. (2018). Fixation reinstatement supports visuospatial memory in older adults. Journal of Experimental Psychology: Human Perception and Performance, 44(7), 1119–1127. https://doi.org/10.1037/xhp0000522

Wynn, J. S., Shen, K., & Ryan, J. D. (2019). Eye Movements Actively Reinstate Spatiotemporal Mnemonic Content. Vision, 3(2), 21. https://doi.org/10.3390/vision3020021

Yassa, M. a, & Stark, C. E. L. (2011). Pattern separation in the hippocampus. Trends in Neurosciences, 34(10), 515–525. https://doi.org/10.1016/j.tins.2011.06.006

Ye, Z., Zhu, B., Zhuang, L., Lu, Z., Chen, C., & Xue, G. (2016). Neural Global Pattern Similarity Underlies True and False Memories. Journal of Neuroscience, 36(25), 6792– 6802. https://doi.org/10.1523/JNEUROSCI.0425-16.2016

